# Both tobacco TA29 and sesame GN13 promoters provide anther-specific GUS expression in transgenic sesame (*Sesamum indicum* L.)

**DOI:** 10.1101/2024.12.11.627888

**Authors:** Anirban Jyoti Debnath, Debabrata Basu, Samir Ranjan Sikdar

## Abstract

Yield improvement is one of the most concerning areas of the “Queen of Oilseed” sesame (*Sesamum indicum* L.) for its successful commercialisation. Heterosis breeding is an alternate approach for improving sesame compared to the time- and labour-consuming conventional breeding. However, tedious hand emasculation and pollination processes restrict the implementation of commercial heterosis on sesame. The unavailability of male sterile, restorer, and maintainer lines further complicates the problem. Biotechnological gene manipulation can silence anther-specific vital gene(s) leading to male sterility. Anther-specific gene study is therefore crucial to reach such a goal. In this study, we have cloned two established anther-specific promoters: sesame SiBGproplus (hereafter GN13, NCBI accession no. *KT246471*) and tobacco TA29 (NCBI accession no. *X52283*) in the plant expression vector pCAMBIA2301 producing respective GN13::GUS and TA29::GUS chimeric vectors. We recovered putatively transgenic T0 sesame lines using conventional *Agrobacterium*-mediated transformation with 1.76% and 1.71% frequencies, respectively. The T0 lines were segregated at the 3:1 Mendelian segregation ratio for kanamycin resistance and generated T1 transgenic lines by self-fertilisation. Most of the T1 transgenics had a single copy of transgene integration. The histological assay revealed the tapetum-specific GUS expression in the T1 transgenic lines; GUS expression was undetected in other tested plant parts. The results depict successful anther/tapetum-specific expression of two promoters in transgenic sesame lines. In further research, these promoters could be used for anther-specific cytotoxic gene expression, RNAi, or CRISPR-mediated mutational approaches to destroy androecium selectively and exhibit male sterility in sesame, resulting in heterosis-mediated improvement.

## Introduction

The exponential human population growth and simultaneous agricultural resource depletion will soon create a food crisis (United Nations 2009, 2022). Moreover, vegetable oil consumption is also rising (Debnath et al. 2024a). This urgent situation demands increased food crop production, including oilseed crops. Despite its excellent oil quality, the “Queen of Oilseed” sesame (*Sesamum indicum* L., family Pedaliaceae) suffers a lower yield problem. The inadequate yield hinders its successful commercialisation (Debnath et al. 2018, 2024a).

Conventional breeding or biotechnological approaches involving gene manipulation can improve sesame yield. However, conventional breeding is a slow process and limited efforts have been taken to increase sesame yield (Debnath et al. 2014; Teklu et al. 2022; Qureshi et al. 2022). Heterosis breeding is an alternative approach to potentially increase sesame yield (Ragiba and Reddy 2000; Mothilal and Ganesan 2005; Yadav et al. 2005; Khan et al. 2009; Prajapati et al. 2009; Shobha Rani et al. 2015; Virani et al. 2017; Ismail et al. 2020; Kumar et al. 2022). Still, commercial heterosis exploitation in sesame is impractical due to labour-intensive hand emasculation and pollination techniques (Yermanos and Osman 1981; Prabakaran 1998). Using sesame male sterile and restorer lines in breeding programs can resolve this issue. However, the unavailability of male sterile, restorer, and maintainer lines primarily constrain sesame heterosis exploitation (Mariani et al. 1990; Carlsson et al. 2008).

Anther-specific gene(s) silencing to inhibit pollen production could alternatively produce genetically induced male sterility (Nawaz-ul-Rehman et al. 2008). Such biotechnologically induced male sterile lines are useful for heterosis breeding. This transgenic “molecular heterosis” approach has been reported in different plants previously (Kultunow et al. 1990; Mariani et al. 1990; Hernould et al. 1993; Rong et al. 1996; Burgess et al. 2002; Tehseen et al. 2010; Jing et al. 2012; Rao et al. 2017; Wan et al. 2019; Singh et al. 2021). However, there is no report to induce male sterility through genetic engineering in sesame to date. Therefore, the anther-specific gene/promoter assessment could lead to genetic male sterile sesame plants in further research.

Previously, we identified a sesame anther-specific promoter named SiBGproplus (hereafter GN13, NCBI accession no. *KT246471*). It is the upstream region of the sesame homolog of tobacco (*Nicotiana tabacum* L.) *ß-1,3-glucanase* gene – a diverse anther/tapetum-specific gene family encoding well-characterised plant hydrolytic enzymes. However, some family members are also expressed in roots, leaves, and other floral tissues (Thimmapuram et al. 2001; Wan et al. 2011). The anther-expressing gene class, callase, executes an essential pollen developmental function of tetrads callose (*β*-1,3-glucan) wall dissolution (Stieglitz 1977). Previously, we characterised the GN13 promoter and demonstrated its anther/tapetum-specific GN13 expression in a transgenic tobacco model (Parveen et al. 2018).

*TA29* is one of the most important genes specifically expressed in the tapetum layer during tobacco anther development (Koltunow et al. 1990; Evrard et al. 1991; McCormick 1991; Scott et al. 1991). The probable roles of *TA29*-encoded membrane-bound 20-kDa protein are membrane transport of nonpolar compounds and involvement in pollen outer wall formation (Pacini et al. 1985; Schrauwen et al. 1996). The TA29 promoter (NCBI accession no. *X52283*) is well-known for its tapetum-specific expression in tobacco and other plants (Koltunow et al. 1990; Mariani et al. 1990; Denis et al. 1993; Kriete et al. 1996; Rosellini et al. 2001; Kavita and Burma 2008; Arun et al. 2011; Madhuri et al. 2012; Nandy et al. 2013).

In this article, we are depicting transgenesis-mediated anther-specific GUS expressions driven by GN13 and TA29 in sesame. This is the first report of anther-specific promoter-derived gene expression in sesame.

The seeds of sesame (*Sesamum indicum* L.) cultivar JK-1 and tobacco (*Nicotiana tabacum* L.) cultivar SR1 were collected from the National Bureau of Plant Genetic Resources, Pusa, New Delhi, India, sown on ½MS medium (Murashige and Skoog 1962) supplemented with 3% sucrose (w/v) and 0.8% agar (w/v), and incubated at 28±2°C in the dark for germination. The germinated seedlings were clonally propagated at 25±2°C under the 16/8 photoperiod using apical shoot tip culture in MS medium and refreshed monthly.

Using an adapter-mediated directional cloning strategy, the sesame 793-bp GN13 and tobacco 871-bp TA29 promoters were cloned to construct recombinant pCAMBIA 2301 plant expression vectors according to Debnath et al. (2024b). Briefly, the sesame GN13 and tobacco TA29 promoters were amplified from the respective clonally propagated leaf tissues using genomic PCR utilising GN13-F/R and TA29-F/R primer pairs (Table 1). The primers contained adapter sequences for the recognition site of *Hind*III and *Nco*I to facilitate downstream directional cloning. The GN13 and TA29 were cloned into the plant expression vector pCAMBIA 2301 by replacing its CaMV35S promoter at *Hind*III and *Nco*I site, fused with the GUS reporter gene, and named GN13::GUS and TA29::GUS, respectively (Fig. 1a, b). The recombinant plasmids had the *Neomycin phosphotransferase II* (*NPT II*) gene conferring kanamycin resistance as the plant selection marker. The clones were analysed and confirmed by PCR, restriction digestion, and DNA sequencing (Table 1). The plasmids from confirmed clones were mobilised into *Agrobacterium tumefaciens* LBA4404 and used in plant transformation experiments.

**Table 1.**
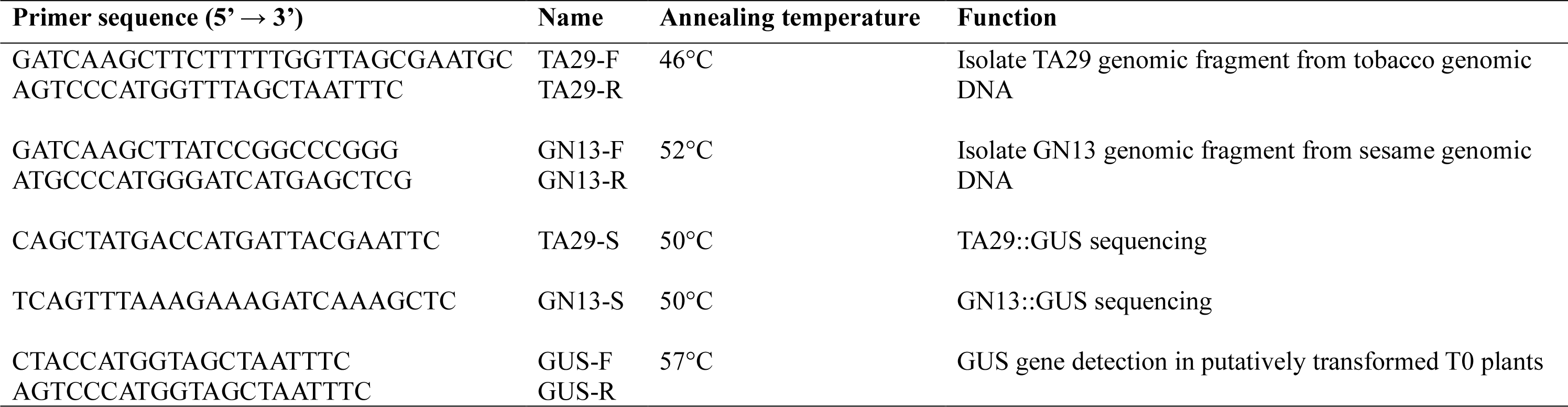
Primers used in this study.

**Fig. 1.**
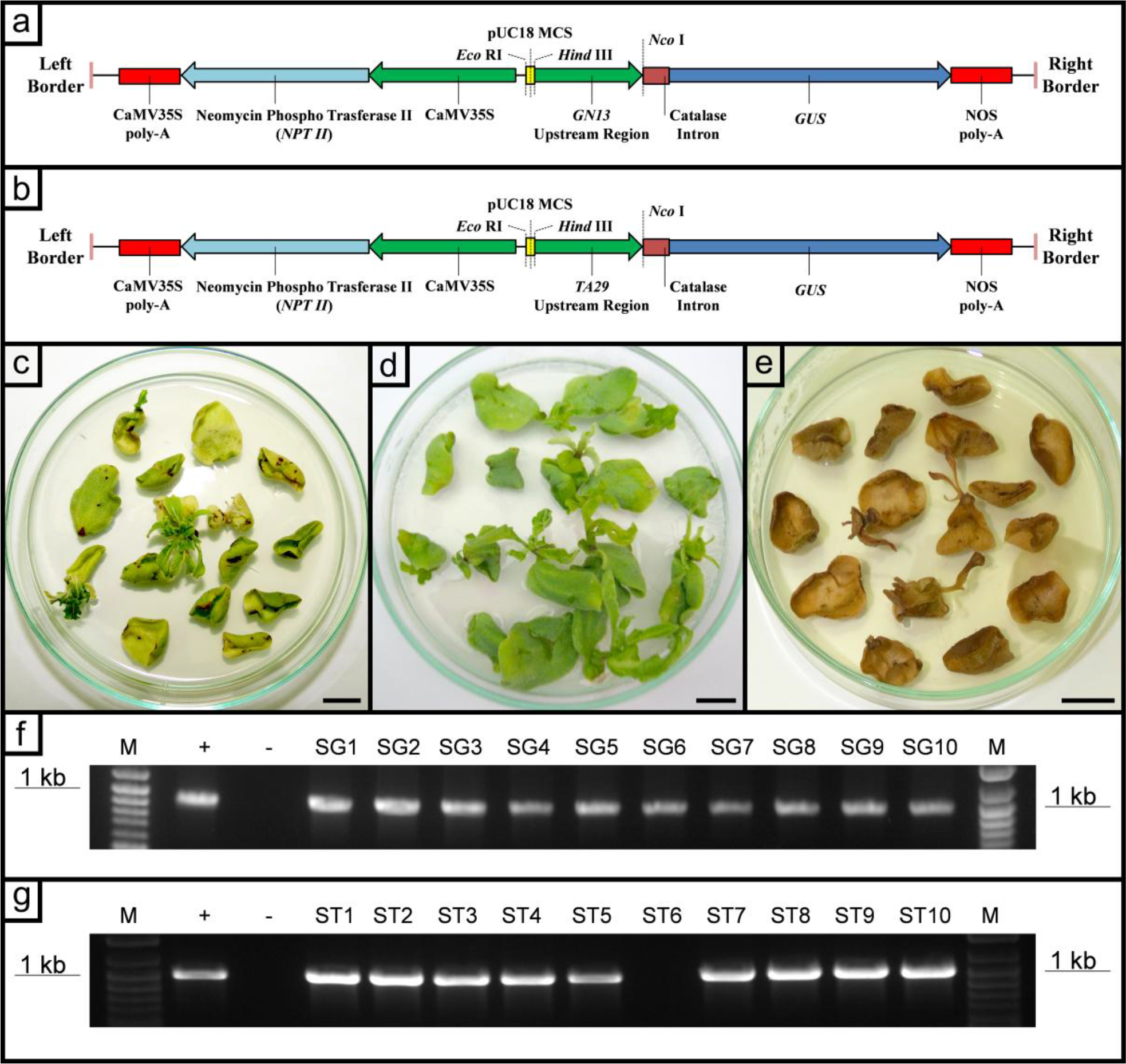
Conventional *Agrobacterium*-mediated sesame transformation with GN13::GUS and TA29::GUS and PCR validation of putative transgenic lines. The T-DNAs of **(a)** GN13::GUS and **(b)** TA29::GUS consist of primarily three components – a CaMV35S::NPTII::CaMV35S poly-A cassette near the left border, a GN13/TA29::Catalase::GUS::NOS poly-A cassette near the right border, and a pUC18 multiple-cloning site (MCS) in between. The GN13/TA29 promoter was cloned into the *Hind*III-*Nco*I site. The unique *Eco*RI site in the MCS is used in the Southern blotting; **(c)** putatively transformed shoots under kanamycin selection; **(d)** non-transformed cotyledons cultured without kanamycin selection; **(e)** non-transformed cotyledons cultured under kanamycin selection; visualisation of GUS bands by PCR-mediated 1% agarose gel electrophoresis in ten randomly selected putatively transformed T0 **(f)** GN13::GUS, and **(g)** TA29::GUS lines

The 4-day-old sesame cotyledons were transformed by GN13::GUS or TA29::GUS adopting the conventional *Agrobacterium*-mediated transformation method described by Yadav et al. (2010) with necessary modifications (Fig. 2). In five experiments, GN13::GUS and TA29::GUS were mobilised in 850 and 820 sesame cotyledon explants, respectively. Shoots were selected using 50 mg/L kanamycin and regenerated according to Debnath et al. (2018) to develop putatively transformed T0 lines (Fig. 1c, d, e). The kanamycin-resistant T0 lines were screened by genomic PCR (95°C, 2 minutes; [95°C, 30 s; 57°C, 45 s; 72°C, 1 minute] 35x; 72°C, 7 minutes; 4°C, infinity) using GUS-F/R primers (Table 1) and the 1.1-kb band of the GUS fragment was visualised by 1% gel electrophoresis (Fig. 1f, g). As a result, 15 and 14 kanamycin-resistant *GUS*-PCR positive putative pGN13::GUS and pTA29::GUS T0 transformed plants were obtained, respectively. The transformation frequency was calculated by the following formula:

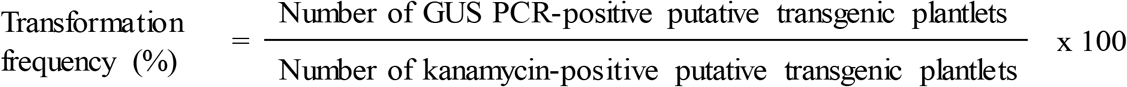

**Fig. 2.**
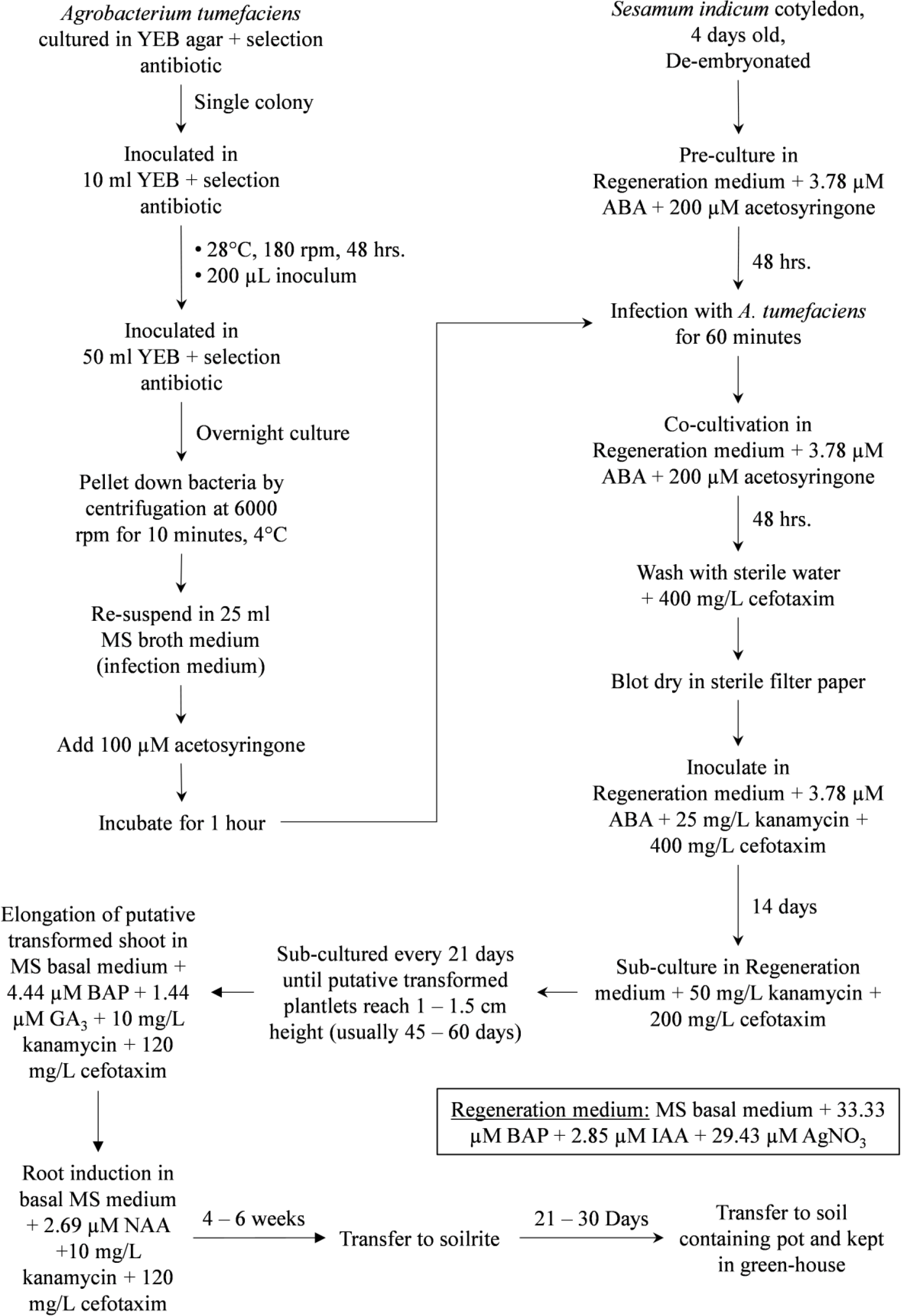
Steps of conventional *Agrobacterium*-mediated transformation of sesame

The average transformation frequencies of 5 experiments of GN13::GUS and TA29::GUS transformation were 1.76% and 1.71%, respectively (Table 2).

**Table 2.**
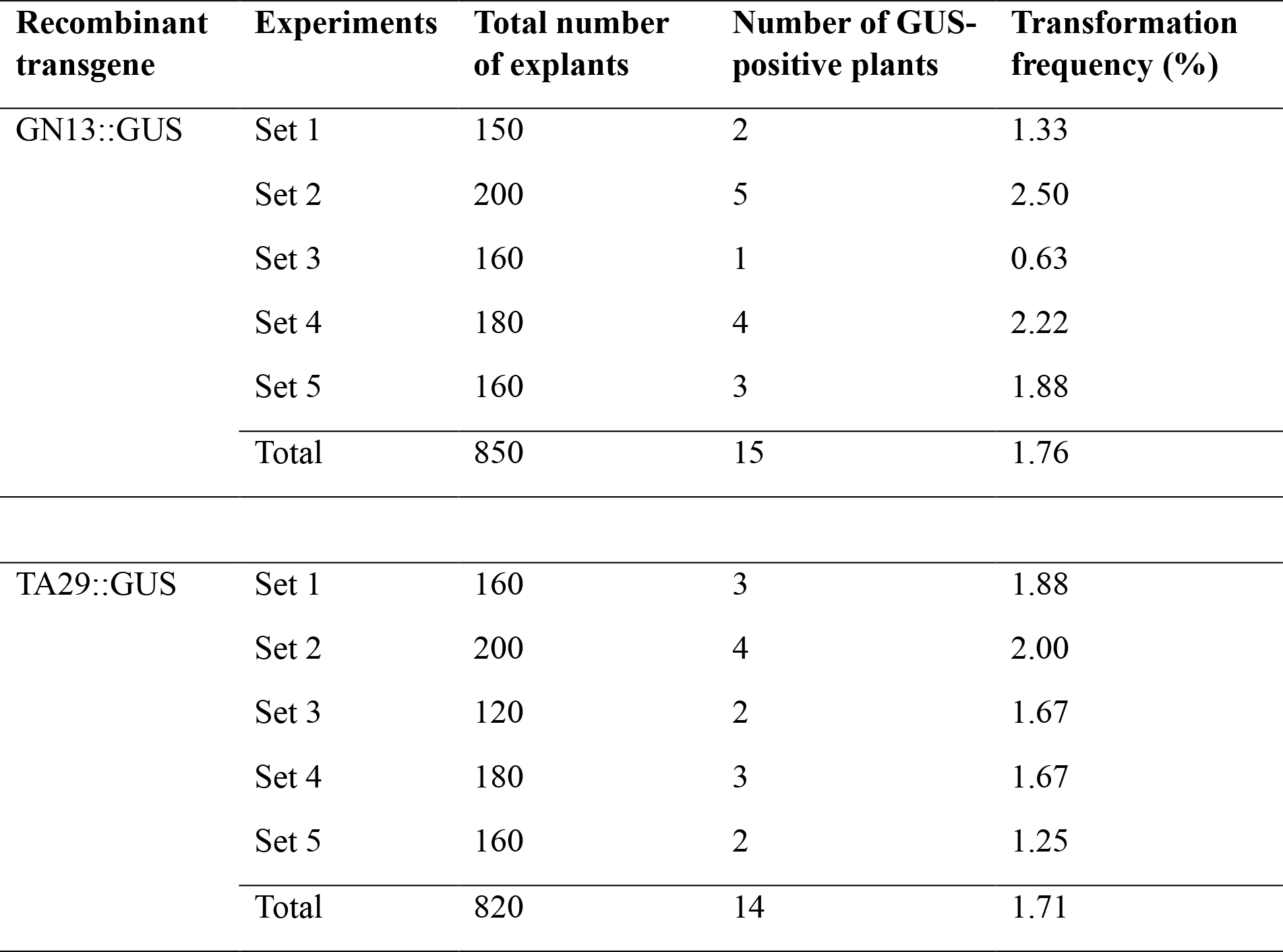
Transformation frequency of sesame, transformed by recombinant GN13::GUS and TA29::GUS by the conventional *Agrobacterium*-mediated transformation method.

All the putatively transformed T0 plants were fertile and produced T1 progenies by self-fertilisation. Ten T0 lines were randomly selected and their seeds were germinated in ½MS medium supplemented with 50 mg/L kanamycin to raise T1 progenies. Segregation analysis on the number of germinated and non-germinated seedlings by χ2 test revealed that the T1 progeny plants followed the 3:1 Mendelian segregation ratio for a single dominant trait of kanamycin resistance (Table 3).

**Table 3.** Mendelian segregation ratio by χ^2^ test of the transformed plants.

| Recombinant transgene | Line No. | Number of tested seeds | Kanamycin<sup>R</sup> | Kanamycin<sup>S</sup> | $\chi^2$ value | Segregation ratio |
| --- | --- | --- | --- | --- | --- | --- |
| GN13::GUS | 1 | 110 | 80 | 30 | 0.303 | 2.67:1 |
|  | 2 | 117 | 87 | 30 | 0.026 | 2.9:1 |
|  | 3 | 101 | 77 | 24 | 0.083 | 3.21:1 |
|  | 4 | 107 | 82 | 25 | 0.153 | 3.28:1 |
|  | 5 | 119 | 88 | 31 | 0.070 | 2.84:1 |
|  | 6 | 115 | 84 | 31 | 0.235 | 2.71:1 |
|  | 7 | 103 | 79 | 24 | 0.159 | 3.29:1 |
|  | 8 | 109 | 83 | 26 | 0.076 | 3.19:1 |
|  | 9 | 114 | 87 | 27 | 0.105 | 3.22:1 |
|  | 10 | 116 | 88 | 28 | 0.046 | 3.14:1 |
| TA29::GUS | 1 | 102 | 75 | 27 | 0.118 | 2.78:1 |
|  | 2 | 121 | 93 | 28 | 0.223 | 3.32:1 |
|  | 3 | 119 | 90 | 29 | 0.025 | 3.1:1 |
|  | 4 | 105 | 77 | 28 | 0.156 | 2.75:1 |
|  | 5 | 110 | 81 | 29 | 0.109 | 2.79:1 |
|  | 7 | 107 | 79 | 28 | 0.078 | 2.82:1 |
|  | 8 | 109 | 80 | 29 | 0.150 | 2.76:1 |
|  | 9 | 114 | 86 | 28 | 0.012 | 3.07:1 |
|  | 10 | 103 | 75 | 28 | 0.262 | 2.68:1 |
|  | 11 | 115 | 89 | 26 | 0.351 | 3.42:1 |

Kanamycin<sup>R</sup> = kanamycin-resistant plants; Kanamycin<sup>S</sup> = kanamycin-sensitive plants

According to Debnath et al. (2024c), the T1 transgenic lines were analysed by Southern blot to confirm the transgene integration and its copy number. The T-DNA region of GN13::GUS and TA29::GUS cassettes contain a single *Eco*RI site at the multiple-cloning site (Fig. 1a, b). Genomic DNAs of the T1 transgenic plants were digested with *Eco*RI and hybridised with the radio-labelled probe prepared against the promoter sequences. We detected a 17-kb band in all Southern blot lanes of GN13::GUS transgenics, including the non-transformed control (arrowhead in Fig. 3a), which signified the presence of a single copy endogenous GN13 promoter in sesame. No such endogenous band was present in the TA29::GUS Southern blot. The presence of two distinct bands of the SG4 line (GN13::GUS transgenic) and a faint second band of the ST1 line (TA29::GUS transgenic) suggested the insertion of two copies of T-DNA in the genomic DNA of those sesame transgenic lines. The rest of the T1 transgenic lines had a single copy of transgene. No hybridisation signal was detected from the non-transformed plants (Fig. 3a, b).

**Fig. 3.**
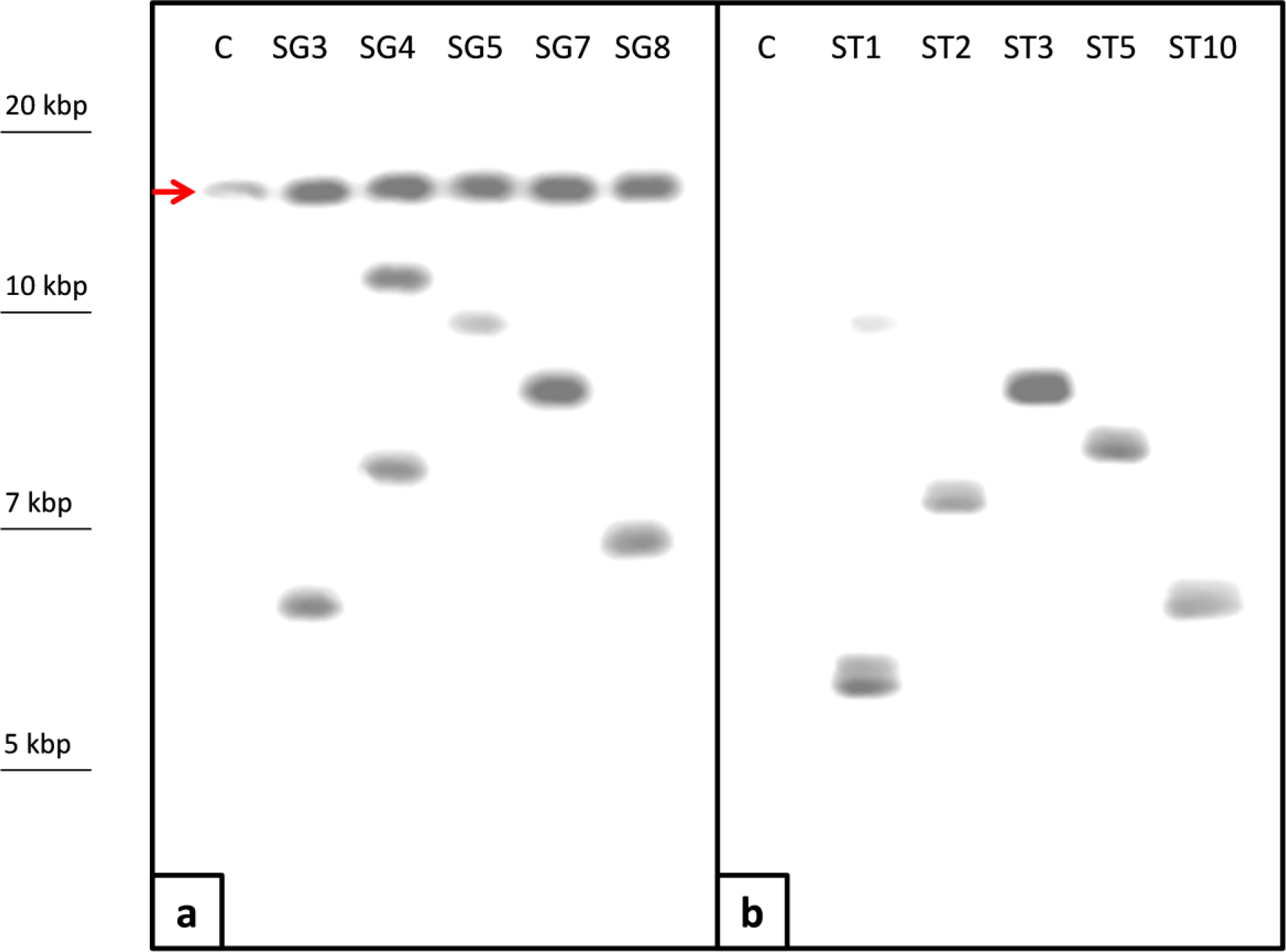
Southern blot hybridisation of five T1 transgenic lines. **(a)** The GN13::GUS T1 transgenics harbour a 17-kb endogenous band of GN13 in all the lanes, including control. The rest of the lanes contain a single band confirming the single copy of transgene integration except the SG4 where the appearance of two bands suggested genomic integration of transgene in two different loci. **(b)** The TA29::GUS T1 transgenics blot lanes contain a single band confirming the single copy of transgene integration except for the ST1 where the second faint band indicated a possible genomic integration of transgene in two different loci

Since the existence and integrity of the tapetum layer are temporally correlated with flower bud size (Koltunow et al. 1990; Mariani et al. 1990), elucidation of the correct bud size containing intact tapetum layer is pivotal in studying tapetum-specific gene expression. Earlier we reported that 4 mm-sized 4-day-old sesame flower buds contain an intact tapetum layer (Fig. 4) (Parveen et al. 2018). Therefore, to monitor the anther/tapetum-specific promoter(s)-driven GUS expression, we performed the histological GUS assay (Jefferson et al. 1987) of 4-day-old T1 transformed and non-transformed anthers. GUS-stained anthers were fixed in FAA (5 ml 37% formalin, 5 ml glacial acetic acid, and 90 ml 85% ethanol) and kept in melting ice for 6 hours. The fixed anthers were sequentially dehydrated in an alcohol gradient series (70%:16 hrs., 80%:30 minutes, 90%:1 hr., 95%:1 hr., and absolute:1 hr.). Dehydrated anthers were passed through a series of the alcohol-chloroform gradient (alcohol:chloroform 3:1, 1:1, and 1:3 sequentially, keeping in each solution for 1 hour) and finally transferred to pure chloroform. Surgipath Paraplast (Leica, Germany, melting point 60°C) chips were added to the chloroform containing the anthers and kept open at 60°C for two days to evaporate the chloroform. Finally, paraplast blocks were prepared from the processed anthers and subsequently sectioned in a sliding microtome (SM2010R, Leica, Germany) at the thickness of 10 μm. Anther sections were collected in 0.1% poly-L-lysine (Sigma-Aldrich, USA) coated glass slides. Collected sections were cleared by chloroform, hydrated by passing them through alcohol gradients (absolute, 95%, 90%, 80%, 70%, 50%, and 30%), kept for five minutes in each solution and finally placed in water. Finally, a drop of D.P.X. Mountant (HiMedia, India) was poured over the section, covered with a cover slip, and observed under a light microscope (Axioshop 40, Carl Zeiss, Germany). Mature flowers without anther, leaf, and root tissues of the non-transgenic and T1 progeny plants were also GUS-stained, subsequently bleached with alcohol, and observed for blue spots.

**Fig. 4.**
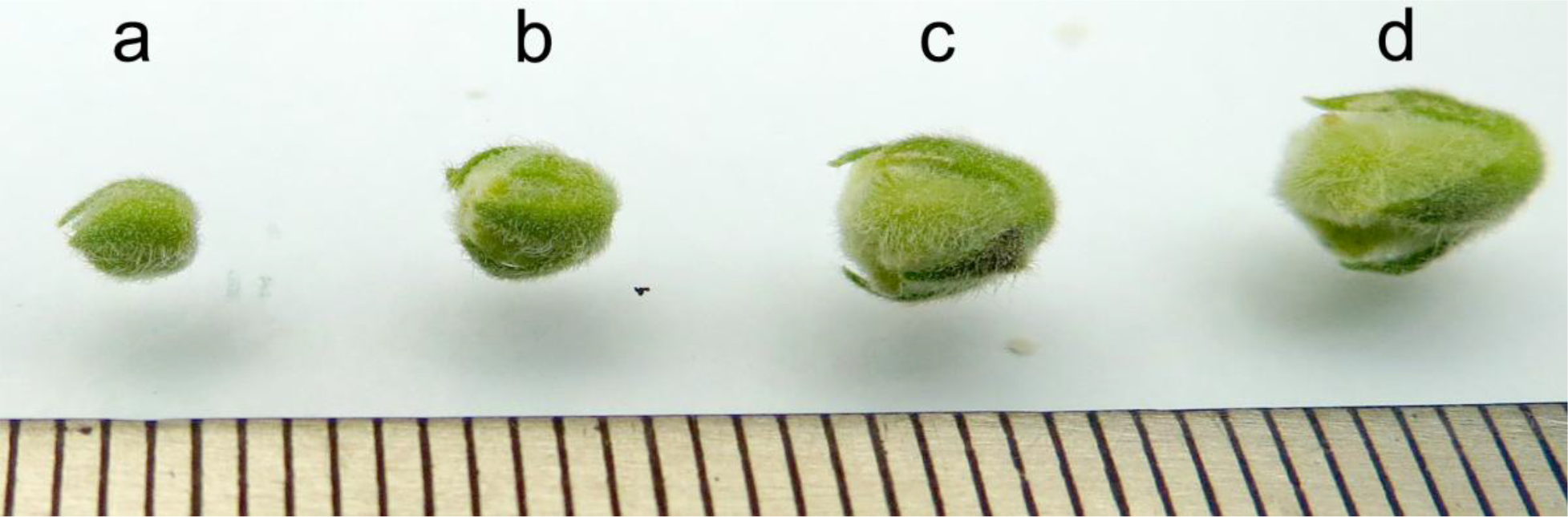
Different-aged sesame flower buds for histological GUS assay; (a) 3-day-old; (b) 4-day-old; (c) 5-day-old; and (d) 6-day-old. Since the tapetum integrity varies with the bud’s age, having a perfect-sized bud with an intact tapetum layer is crucial from anther-specific GUS expression analysis. We used the 4-day-old bud for GUS assay since in an earlier study we found anthers from buds of this age contained intact tapetum layer and sporogenous tissue (Parveen et al. 2018). The bottom of the image displays a millimetre scale

The GN13 is the promoter of the sesame homologue of tobacco *ß-1,3-glucanase*. Previously, we demonstrated its anther-specific expression in tobacco (Parveen et al. 2018). In conjunction, we showed GN13-mediated homologous GUS expression in the anther/tapetum layer of the T1 transgenic sesame in this study (Fig. 5b); other plant organs had undetectable GUS activity (Fig. 5e, h, k). Several studies reported the anther-specific expression of *ß-1,3-glucanase* (Perrot et al. 2022; Pei et al. 2023; Zhang et al. 2024) and its expression manipulation could lead to male sterility in crop and model plants (Frankel et al. 1969; Warmke and Overman 1972; Hird et al. 1993; Jin et al. 1997; Worrall et al. 1992; Tsuchiya et al. 1995; Wan et al. 2011; Liu et al. 2014).

**Fig. 5.**
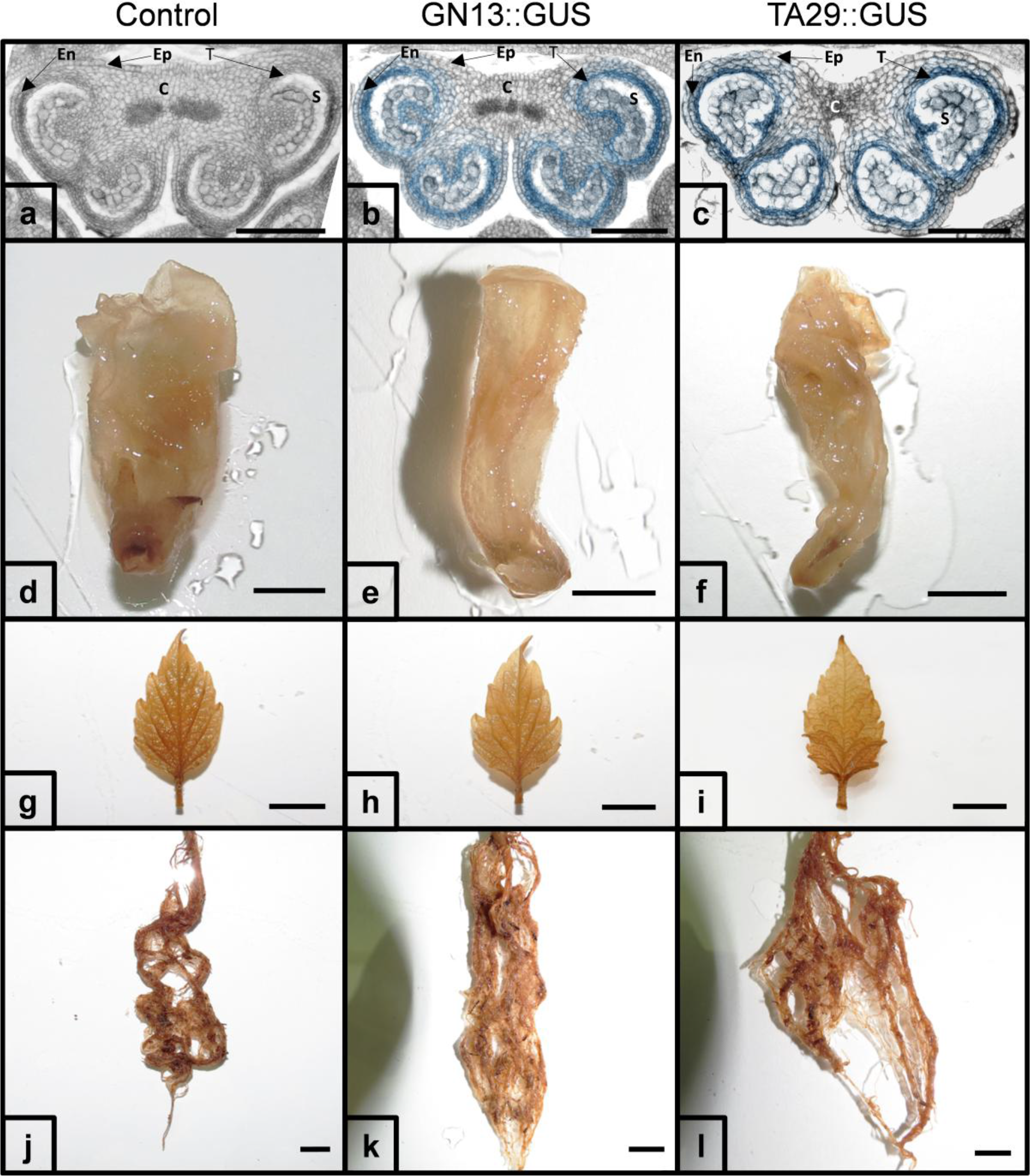
Histological GUS assay of anther/tapetum-specific promoter expression. **(a)** The GUS expression was not detected in the non-transformed control anther section; the tapetum layers of **(b)** GN13::GUS and (C) TA29::GUS T1 transgenic anther sections stained blue confirming the ability of anther/tapetum-specific expression of GN13 and TA29 promoters in transgenic sesame; the blue spots of GUS were undetected in **(d to f)** flowers without anthers, **(g to i)** leaves, and **(j to l)** roots of the non-transformed and transformed plants. *C* connective, *En* endothecium, *Ep* epidermis, *S* sporogenous tissue, *T* tapetum

The anther/tapetum-specific *TA29* promoter expression has been well-established in tobacco (Koltunow et al. 1990; Mariani et al. 1990; Kriete et al. 1996; Arun et al. 2011; Madhuri et al. 2012). In this study, we found TA29-mediated anther/tapetum layer-exclusive GUS expression in the heterologous system of sesame T1 transgenics (Fig. 5c) and it was undetected in other transgenic plant parts (Fig. 5f, i, l). Consistent with our findings, the *TA29* promoter is also reported to be expressed in other heterologous plant systems such as *Brassica napus* (Denis et al. 1993), *Brassica juncea* (Kavita and Burma 2008), alfalfa (Rosellini et al. 2001), cotton (Soni et al. 2021), and tomato (Nandy et al. 2013; Jogam et al. 2024).

Promoters are crucial *cis*-acting elements that regulate gene expression transcriptionally. Tissue-specific promoters are specific promoter types that spatially regulate gene expression. Several reports on the tissue-specific promoter expression study provided key gene regulation and plant developmental information for plant biotechnological improvements (Shah et al. 2015; Kummari et al. 2020; Singha et al. 2021; Danila et al. 2022; Zhao et al. 2022; Brooks et al. 2023). Researchers have emphasised the importance of anther-specific expression studies in furnishing essential details of spatiotemporal gene regulation cues useful for further plant breeding programs (Luo et al. 2006; Gupta et al. 2007; Roque et al. 2019; Shi et al. 2021; Zhao et al. 2022; Lei et al. 2023; Szeluga et al. 2023; Li et al. 2024). In this study, we have depicted anther-specific expression of two promoters, GN13 and TA29 in transgenic sesame. In future research, these promoters could be adopted for anther-specific cytotoxic gene expression, RNAi, or CRISPR-mediated mutational approach (Debnath et al. 2024a) to selectively destroy androecium and manifest male sterility in sesame leading to heterosis-mediated improvement.

## Abbreviations

CRISPR: Clustered regularly interspaced short palindromic repeats
FAA: Formaldehyde Alcohol Acetic Acid
GUS: β-glucuronidase
MS: Murashige and Skoog
NPT: Neomycin phosphotransferase
PCR: Polymerase chain reaction
RNAi: RNA interference

## Acknowledgements

This work was supported by the National Agricultural Innovation Project, Indian Council of Agricultural Research (ICAR-NAIP) under Grant [Funding Number: NAIP/C4/C1090].

## Author contribution

Debabrata Basu and Samir Ranjan Sikdar arranged the funds and resources for the research, designed the experiments, and supervised the work; Anirban Jyoti Debnath performed the experiments, analysed the data, prepared the tables, graphics, and first draft of the manuscript; all the authors revised and approved the final manuscript.

## Declarations

### Statements

The authors disclose no financial or non-financial interests directly or indirectly related to the work submitted for publication. They also have no relevant competing interests to declare to the content of this article.

